# Natural Variation in *Drosophila melanogaster Clock* PolyQ Length: Geographic Gradients and Functional Properties

**DOI:** 10.64898/2026.02.03.703442

**Authors:** Maya Yair, Bettina Fishman, Martin Kapun, Eran Tauber

**Affiliations:** Department of Evolutionary and Environmental Biology, and Institute of Evolution, University of Haifa 199 Abba-Hushi Avenue, Haifa 3498838 Israel; Naturhistorisches Museum Wien, Burgring 7, 1010 Wien, Austria

**Keywords:** *Clock* gene, circadian rhythm, polyglutamine polymorphism, temperature compensation, climatic adaptation, *Drosophila melanogaster*, natural variation, geographic cline

## Abstract

Natural variation in circadian clock genes provides a powerful framework for understanding how organisms respond to environmental heterogeneity. The *Clock* (*Clk*) gene encodes a core transcriptional regulator of circadian rhythms and contains a polymorphic polyglutamine (polyQ) tract whose evolutionary significance remains unclear. Here, we integrate population genomic, behavioral, and molecular analyses to investigate the functional and geographic patterns of *Clk* polyQ variation in *Drosophila melanogaster*. Using data from 127 European populations, we identify 11 *Clk* polyQ alleles and find that their frequencies are geographically structured, most consistently along an east–west gradient: the Q25 and Q27 alleles show robust clines in longitude and in a bioclimatic axis of continentality, whereas latitudinal and altitudinal trends are weaker. Behavioral assays of near-isogenic lines revealed that polyQ length modulates circadian function under thermal challenge: most alleles maintained stable free-running periods across temperatures, whereas the intermediate-length Q25 allele showed the strongest, though modest, temperature sensitivity. Circadian phase showed pronounced allele-specific sensitivity to elevated temperature in laboratory assays, although phase variation did not display a consistent relationship with geographic variables. At the molecular level, luciferase reporter assays showed that longer polyQ alleles exhibited higher transcriptional activity, linking polyQ length to CLK-mediated gene expression. Together, these results demonstrate that natural variation in *Clk* polyQ length has measurable functional consequences for circadian regulation and is geographically structured in patterns consistent with underlying climatic variation, highlighting the potential for low-complexity regions to modulate clock function in a context-dependent manner across environmental gradients.

## **1** Introduction

Circadian clocks are endogenous timekeeping systems that synchronize physiology and behavior with the 24-hour environmental cycle. They confer a critical adaptive advantage by aligning internal biological processes such as metabolism, reproduction, and locomotor activity with predictable daily changes in light and temperature. Across taxa, disruption of circadian organization reduces fitness through impaired survival, fecundity, and competitive performance (reviewed by Jabbur et al. 2024). The circadian system therefore serves as a key interface between genetic regulation and environmental periodicity, allowing organisms to anticipate rather than simply react to environmental fluctuations.

At the core of the circadian system lies *Clock (Clk)*, a transcription factor that functions as a master regulator of rhythmic gene expression. In *Drosophila melanogaster*, CLK forms a heterodimer with CYCLE (CYC) to activate downstream clock genes, including *period (per)* and *timeless (tim)*, through E-box enhancer elements (Darlington et al., 1998; Patke et al., 2020). The resulting feedback loops involving PER and TIM proteins generate near 24-hour oscillations that persist under constant conditions (Vosshall et al., 1994; Price et al., 1998). CLK activity is further modulated by additional feedback components such as *vrille (vri)*, *pdp1*, and *clockwork orange (cwo)*, which fine-tune transcriptional amplitude and phase (Damulewicz & Mazzotta, 2025). Because CLK sits at the apex of this transcriptional hierarchy, variation in its structure or activity has the potential to influence the entire circadian network.

Circadian rhythms are not rigidly fixed but can evolve in response to environmental pressures. Two key properties, entrainment to external cues and temperature compensation, enable organisms to maintain synchrony with local conditions. In natural populations, clock genes frequently exhibit molecular polymorphisms that affect these traits, suggesting that variation in clock components contributes to local adaptation (Bell-Pedersen et al., 2005; Khatib et al., 2023). For example, *Drosophila* populations across Europe show subtle but significant geographic structure in allele frequencies of several circadian genes, with some variants linked to differences in free-running period, activity phase, and seasonal traits such as diapause and cold tolerance (Khatib et al., 2023). These patterns indicate that natural selection can act on circadian regulatory variation.

Among the core clock components, *Clk* stands out as a particularly promising locus for adaptive evolution. The gene contains a repetitive polyglutamine (polyQ) region in its C-terminal domain, where variation in repeat length has been observed across multiple species. This region influences protein–protein interactions and transcriptional activity, suggesting that natural variation in *Clk* polyQ length could modulate circadian properties such as temperature compensation.

Natural variation in circadian clock genes has been widely documented and often exhibits geographic patterns consistent with local adaptation. A well-studied example involves a repetitive region in the *per* gene that encodes pairs of Threonine–Glycine (Thr–Gly) residues. The most common alleles, (Thr–Gly)_17_ and (Thr–Gly)_20_, show a clear latitudinal cline in European populations of *Drosophila melanogaster*, with the longer repeat more frequent at higher latitudes and a similar, although weaker, trend observed in Australia (Costa et al., 1992; Sawyer et al., 2006). Behavioral studies revealed that flies carrying the (Thr–Gly)_20_ allele exhibit more effective temperature compensation, whereas the (Thr–Gly)_17_ variant shortens the free-running period at low temperatures (Sawyer et al., 1997). These differences suggest that thermal selection maintains allele frequency gradients, favoring variants that optimize circadian performance under local climatic conditions (Hut et al., 2013; Kyriacou et al., 2008).

Similar geographic trends have been identified in *Clk*, highlighting its potential role in adaptive modulation of circadian timing. The *Clk* protein contains a repetitive polyglutamine (polyQ) tract located in its C-terminal glutamine-rich domain, encoded by a tandem array of CAG/CAA codons. Variation in polyQ length has been observed in several taxa and is often associated with environmental gradients. In Chinook salmon (*Oncorhynchus tshawytscha*), for instance, eight distinct *Clk* polyQ alleles were identified, with shorter alleles predominating in southern, low-latitude populations and longer alleles more common at higher latitudes (O’Malley & Banks, 2008). This variation correlates with migration timing, as individuals carrying longer alleles tend to migrate later in the season, suggesting that polyQ length diversity contributes to photoperiodic adaptation.

A similarly patterned relationship has recently been reported across a much broader taxonomic sample of coral reef fish. A comparative analysis of *Clk* polyQ and Qrich repeat length across 20 damselfish species (Pomacentridae) found a significant positive correlation between combined repeat length and pelagic larval duration, a key life-history trait governing the developmental window before reef settlement. This relationship held at the interspecific level but not within a single species once local habitat availability was accounted for, suggesting that polyQ length may track developmental timing across lineages while additional ecological factors modulate its expression within populations (Schalm et al., 2025).

Comparable *Clk* polymorphisms have also been described in avian species. In the bluethroat (*Luscinia svecica*) and the blue tit (*Cyanistes caeruleus*), seven and nine polyQ alleles were identified, respectively. In blue tits, allele length showed a positive correlation with breeding latitude, whereas no such relationship was observed in the migratory bluethroat, possibly due to differences in reliance on local photoperiodic cues (Johnsen et al., 2007). These studies together support the idea that *Clk* polyQ length variation may provide a molecular basis for latitude-dependent adjustments in circadian or seasonal timing.

The functional roles of polyQ tracts further reinforce this interpretation. PolyQ regions are common in transcriptional regulators and can modulate protein–protein interactions, structural flexibility, and transcriptional activity (Gemayel et al., 2015; Mier & Andrade-Navarro, 2021). Moderate variation in polyQ length can tune transcriptional output, while extreme expansions beyond a critical threshold can lead to pathological aggregation (Lieberman et al., 2018). Experimental evidence from plants such as *Arabidopsis thaliana* and *Populus* shows that polyQ variation influences thermal responsiveness and protein localization (Sureshkumar et al., 2025). These findings suggest that naturally occurring variation in *Clk* polyQ length could serve as a mechanism for fine-tuning transcriptional regulation and temperature compensation within the circadian system.

Despite extensive knowledge of the circadian molecular network, the contribution of *Clk* polymorphism to adaptive regulation of clock function remains unclear. The presence of a variable polyQ tract in *Clk*, coupled with its central regulatory role, suggests a plausible mechanism through which natural selection could fine-tune circadian timing and temperature compensation. This study aims to test whether variation in *Clk* polyQ length influences circadian performance under different thermal conditions and to evaluate its potential role in local adaptation.

## **2** Methods and Materials

### 2.1 Analysis of *Clk* polyQ Allele Frequency Across Europe

A large genomic dataset of wild *Drosophila melanogaster* populations across Europe was obtained from the DrosEU consortium (Kapun et al., 2020). DNA sequences from 169 wild populations, each sequenced using Pool-seq, were assembled with the SAMtools package (Li et al., 2009). Reads spanning the *Clk* polyQ region (Chr3L:7,764,779–7,764,908) were extracted together with flanking indel polymorphisms. Known indel variants corresponding to distinct polyQ alleles were manually counted by visual inspection of read alignments in the Integrative Genomics Viewer (IGV; Robinson et al., 2011). Allele frequencies were calculated by dividing the number of reads supporting each indel by the total read depth at that site.

To assess environmental associations, a principal component analysis (PCA) was conducted using 19 bioclimatic variables obtained from the WorldClim database (https://worldclim.org; Fick & Hijmans, 2017). The variables included annual means (for example, temperature and precipitation), measures of seasonality (for example, annual ranges in temperature and precipitation), and extreme environmental values (for example, temperature of the coldest and warmest months and precipitation of the wettest and driest quarters).

To test for associations between *Clk* polyQ allele frequencies and geographic or climatic variables, we used generalized linear models (GLMs) implemented in R (RStudio version 2025.05.1). Allele frequencies (proportion of reads supporting a given allele) were modelled with a quasibinomial error and a logit link, which accommodates the overdispersion of Pool-seq allele counts relative to the binomial expectation. For each allele, one model included latitude, altitude and longitude as joint predictors, and a separate model included the first two climatic principal components (PC1 and PC2) derived from the bioclimatic PCA as explanatory variables. To avoid conflating spatial with seasonal variation, the geographic and climatic association analyses were restricted to spring-collected samples (101 of the 127 populations). Because DrosEU populations were sampled in different seasons and allele frequencies at clock loci can shift seasonally (Bergland et al. 2014), analysing a single season isolates the spatial (clinal) signal from seasonal change. Effect sizes were expressed as odds ratios (OR), which describe how the odds of observing a given allele change with increasing values of the predictor variable (OR > 1 indicates a positive association, OR < 1 a negative association). Model significance was assessed using Wald tests, and p-values were adjusted for multiple comparisons where appropriate.

### 2.2 Generation of Clk polyQ Allele Series

To establish populations homozygous for specific *Clk* polyQ alleles, virgin females were collected from several DrosEU isofemale lines. Each virgin female was paired with a male from the same isofemale line. DNA from the parental flies was extracted using a single-fly squashing protocol and sequenced to determine their *Clk* polyQ genotype. Only F1 progeny from parental pairs that were homozygous for the same allele were retained. This procedure produced six homozygous populations carrying polyQ alleles Q21, Q25, Q29, Q30, Q31, and Q33. Sequence alignment confirmed that these alleles differed only in the number of consecutive glutamine residues within the uninterrupted polyQ stretch (Figure 1).

**Figure 1.**
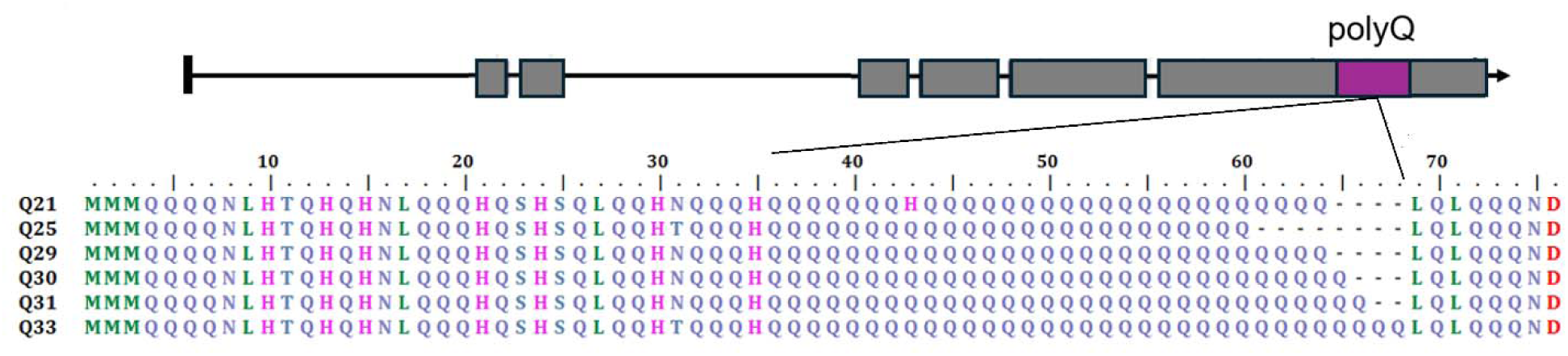
Alignment of six *Clk* polyQ variants used in this study. Each sequence is labeled with “Q” followed by the number of consecutive glutamine residues within the uninterrupted polyQ stretch.

### 2.3 Generation of Near-Isogenic Lines

Near-isogenic lines (NILs) are strains that are genetically identical except for one or a few loci of interest, which minimizes the effects of background genetic variation on phenotypes (Yuan et al., 2017). These lines were produced through a series of backcrosses in which the *Clk* locus from a donor parent was introgressed into the background of a deficiency line, with selection for the desired genotype at each generation (Kooke et al., 2012).

To generate *Clk* polyQ NILs, virgin females from each homozygous *Clk* polyQ line (Q21, Q25, Q29, Q30, Q31, and Q33) were crossed with males carrying a deficiency spanning the *Clk* locus (Df(3L)pbl-X1/TM6C, Sb[1]; BDSC #1420). Wild-type bristled hemizygous females from the F1 offspring were selected and backcrossed with males from the same deficiency stock. This process was repeated for four backcross generations (BC1–BC4). By the fourth generation, progeny were expected to share approximately 96 percent of their genome with the deficiency background, differing primarily at the *Clk* locus. Hemizygous BC4 males were collected for behavioral testing, while females were used for continued backcrossing until at least 20–30 flies per allele were obtained for behavioral assays. The crossing design used to generate the NILs is illustrated in Supplementary Figure S1.

### 2.4 Analysis of Circadian Behavior

Circadian activity was monitored using the Drosophila Activity Monitoring (DAM) system (Trikinetics, Waltham, MA). Flies were first entrained under a 12:12 h light–dark (LD) cycle for five days, followed by nine days in constant darkness (DD). Temperature compensation was examined by testing flies at 24 °C and at 18 °C. Recordings at 29 °C were also attempted, but flies of these near-isogenic lines showed high mortality and loss of measurable rhythmic activity at this temperature and did not yield usable data.

Activity data were analyzed using the BioDare2 webserver (Zielinski et al., 2014) with the Maximum Entropy Spectral Analysis (MESA) method to determine each fly’s free-running period (FRP) that is, the intrinsic circadian period measured under constant conditions without light entrainment. Flies were excluded from analysis if they (i) died before completing at least 10 days (5 LD + 5 DD), (ii) showed arrhythmic behavior, or (iii) displayed FRP values outside the range of 22–28 h. These estimates were flagged by BioDare2 as requiring attention and correspond to weakly rhythmic or otherwise unreliable fits rather than to reliably estimated circadian periods. The criterion was applied uniformly to all genotypes and both temperatures without reference to allele identity (see per-allele counts in Table S4).

Circadian phase was quantified from the entrained 12:12 LD recordings using a single deterministic pipeline written in base R (v4.3.3). Activity was recorded in 30-min bins (48 bins per 24 h). Because the DAM monitors were started at lights-off (ZT12, ∼20:00), each folded profile was rotated so that bin 1 corresponded to ZT0 (lights-on) before any phase reference was computed. For each fly, the LD recording days were folded into an average 24-h activity profile (NA-aware mean per bin). Flies were retained if they showed a mean activity of at least 2 counts per bin over the folding window and non-zero activity on the final day (i.e., alive at the end of recording); only the six established NIL alleles (Q21, Q25, Q29, Q30, Q31, Q33) were included. Because this criterion selects for a scorable LD activity profile rather than for a clean free-running rhythm in DD, the phase dataset (n = 352 at 24 °C; n = 396 at 18 °C) is larger than, and analysed separately from, the FRP dataset described above.

Seven phase references were pre-specified per fly. Within a morning window (ZT21–06) and an evening window (ZT07–16), we extracted the onset, peak, and offset of each activity component, together with the morning–evening (M–E) phase angle, defined as the interval between the morning and evening peaks. Onset and offset were taken as the half-maximum up- and down-crossings of the component (linearly interpolated between bins); the peak was estimated as the centroid (centre of mass above baseline) rather than the single highest bin, giving a more robust estimate for broad or bimodal peaks. Morning references were expressed relative to ZT0 (values after midnight rendered as negative) and the M–E angle as E_peak − M_peak (mod 24 h). The evening references fall predominantly in the anticipatory phase preceding lights-off, limiting the contribution of the light-off startle response.

Genotype effects were tested within each temperature by one-way ANOVA and Kruskal–Wallis on each reference, with Benjamini–Hochberg (BH) correction applied across exactly the seven pre-specified references; Tukey HSD was used for post-hoc pairwise comparisons on references significant after correction. Genotype × temperature effects were tested on the combined dataset by two-way ANOVA, with η² reported as the effect size for each term and the interaction BH-corrected across references and additionally verified by a distribution-free Freedman–Lane permutation test (2000 permutations). For each reference we report the mean resultant length (R□) as a diagnostic of phase concentration; because all references were strongly concentrated (R□ ≥ 0.84, and ≥ 0.98 for the evening peak) and lie away from the ZT0 boundary, phases were summarised with linear statistics. A circular Watson–Williams test gave results concordant with the parametric analysis for all references. All statistics were computed in base R.

### 2.5 Luciferase Assay Constructs and Transformation

Two reporter plasmids were used for the luciferase assay: a *Renilla reniformis* luciferase reporter driven by the copia promoter (Renilla luciferase) for normalization, and a *Drosophila per* E-box firefly luciferase reporter (pGL3-4E-hsp-luc) containing four tandem *per* E-box sequences (CACGTG) with 18 bp of flanking sequence upstream of a minimal *hsp70* promoter. These constructs were generously provided by Sebastian Kadener and Lin Zhang.

The *Clk* expression vector pAC-dCLK was generated by cloning *Clk* cDNA into the pBS-SK(−) vector, followed by subcloning into the NotI site of the pAc5.1-V5/His vector (Invitrogen) for expression in S2 cells (McDonald et al., 2001). The empty pAc5.1-V5/His vector was used as a negative control. All plasmids contained an ampicillin resistance gene for selection.

*Escherichia coli* DH5α cells were transformed using the standard heat-shock protocol. Plasmid DNA was purified with the NucleoSpin Plasmid EasyPure Mini Kit (MACHEREY-NAGEL). The *Clk* polyQ allele in the pAC-dCLK plasmid was verified by Sanger sequencing using primers flanking the polyQ region (pAc-clkL5: AATGGCTGGAAATGCGTGTC; pAc-clkL3: GCTCCGAGTTCAGTGCAAAT; pAc-clkR5: CAGCAACATCAGAGCCACTC; pAc-clkR3: TGCTGCAGGATTTGTTGTTGT). Sequence assembly and analysis were performed in BioEdit, confirming that the original plasmid carried the polyQ33 allele (designated pAc-polyQ33).

Allele sequences from DrosEU isofemale lines were used to generate additional *Clk* expression plasmids carrying polyQ31, polyQ28, and polyQ21 variants. Five µg of pAC-dCLK plasmid DNA and the sequence information for each variant were submitted to GenScript for plasmid construction. The polyQ33 and polyQ31 alleles were selected for analysis due to their high frequencies in the DrosEU dataset, while polyQ28 and polyQ21 were included for their distinct clustering patterns in PCA and unique sequence features, including an internal histidine residue in the Q21 variant. All constructs were verified by Sanger sequencing using the primer sets described above.

### 2.6 S2 Cell Culture, transfection and Luciferase assay

Drosophila Schneider 2 (S2) cells, which lack expression of most endogenous clock genes except *cyc*, were used to evaluate the transcriptional activity of *Clk* polyQ variants. Cells were co-transfected with pAc-*Clk* constructs (polyQ33, polyQ31, polyQ28, or polyQ21) together with the pGL3-4E-hsp-luc reporter plasmid and the Renilla luciferase plasmid for normalization. Cells transfected with the empty pAc5.1-V5/His vector served as negative controls. Luciferase activity was measured 48 hours after transfection.

Luciferase activity was measured to assess the transcriptional regulation of the *Drosophila per* E-box by CLK variants containing different polyQ alleles. Measurements were performed using the Dual-Luciferase Reporter Assay System (Promega) on a BioTek Synergy HTX plate reader with Gen5 software, following the manufacturer’s protocol. Firefly luminescence driven by the pGL3-4E-hsp-luc reporter was measured first, then quenched before measuring Renilla luminescence from the copia-driven reporter (pCopia-Renilla). Relative luciferase activity for each replicate was calculated as the ratio of firefly to Renilla luminescence.

## **3** Results

### 3.1 Spatial Analysis of *Clk* polyQ Allele Frequency Across Europe

Genomic data from 169 *Drosophila melanogaster* populations obtained from the DrosEU dataset (Kapun et al., 2020) were analyzed. After excluding populations with low read coverage and those from extreme latitudes, longitudes, or altitudes, 127 populations remained for analysis. Retained populations spanned latitude 35.1 to 62.6 °N, longitude −8.4 to 38.7 °E, and altitude 11 to 870 m. Within the *Clk* polyQ region, 11 distinct length variants were identified, ranging from Q21 to Q33. To confirm these variants at the sequence level, we extracted the polyQ tract directly from reads spanning the *Clk* repeat across all populations and reconstructed a consensus sequence for each length class (Supplementary Figure S2). The eleven alleles form a nested length series differing only in the number of glutamine codons within the tract, with the longest allele (Q33) recovered from nearly all populations. Within the tract, glutamine is encoded by an interspersed mixture of CAG and CAA codons, and a histidine codon is present near the tract margin in most alleles. The two rarest variants, Q21 and Q32, were each supported by only a small number of reads.

The most common alleles were Q33 and Q27, which occurred in nearly all populations (127 and 122 populations, respectively). Alleles Q25 and Q31 were also frequent, appearing in over 100 populations each. The Q28 allele was moderately common (70 populations), while Q29 and the remaining shorter or longer variants (Q21, Q24, Q26, Q30, Q32) were relatively rare, detected in fewer than 30 populations each.

PolyQ length for each read was determined by calculating the number of base pairs deleted relative to the longest allele, Q33, which consists of 99 base pairs and contains no indels. Structural inspection revealed variation in the positions of insertions and deletions within the polyQ region, indicating that indel events occurred independently across populations. In some alleles, deletions formed a single continuous gap, while in others they appeared as multiple smaller deletions with equivalent total lengths. The Q27 allele showed the greatest diversity of indel configurations, with ten distinct patterns identified. Because the goal of this study was to examine the effect of polyQ length variation rather than indel structure, subsequent analyses focused only on allele length.

To summarize variation in *Clk* polyQ allele composition across Europe, we first performed a principal component analysis (PCA) of allele frequencies for the five most common variants (Q33, Q31, Q28, Q27, and Q25). The first two components explained 58.1% of the total variance (Figure 2; Supplementary Table S1). Populations were distributed continuously along the first two axes, indicating gradual changes in allele composition rather than discrete clusters. PC2 was driven mainly by variation in the frequency of Q28, suggesting that this allele contributes disproportionately to among-population differentiation. These broad patterns motivated subsequent analyses testing the associations between allele frequencies and geographic and climatic variables.

**Figure 2.**
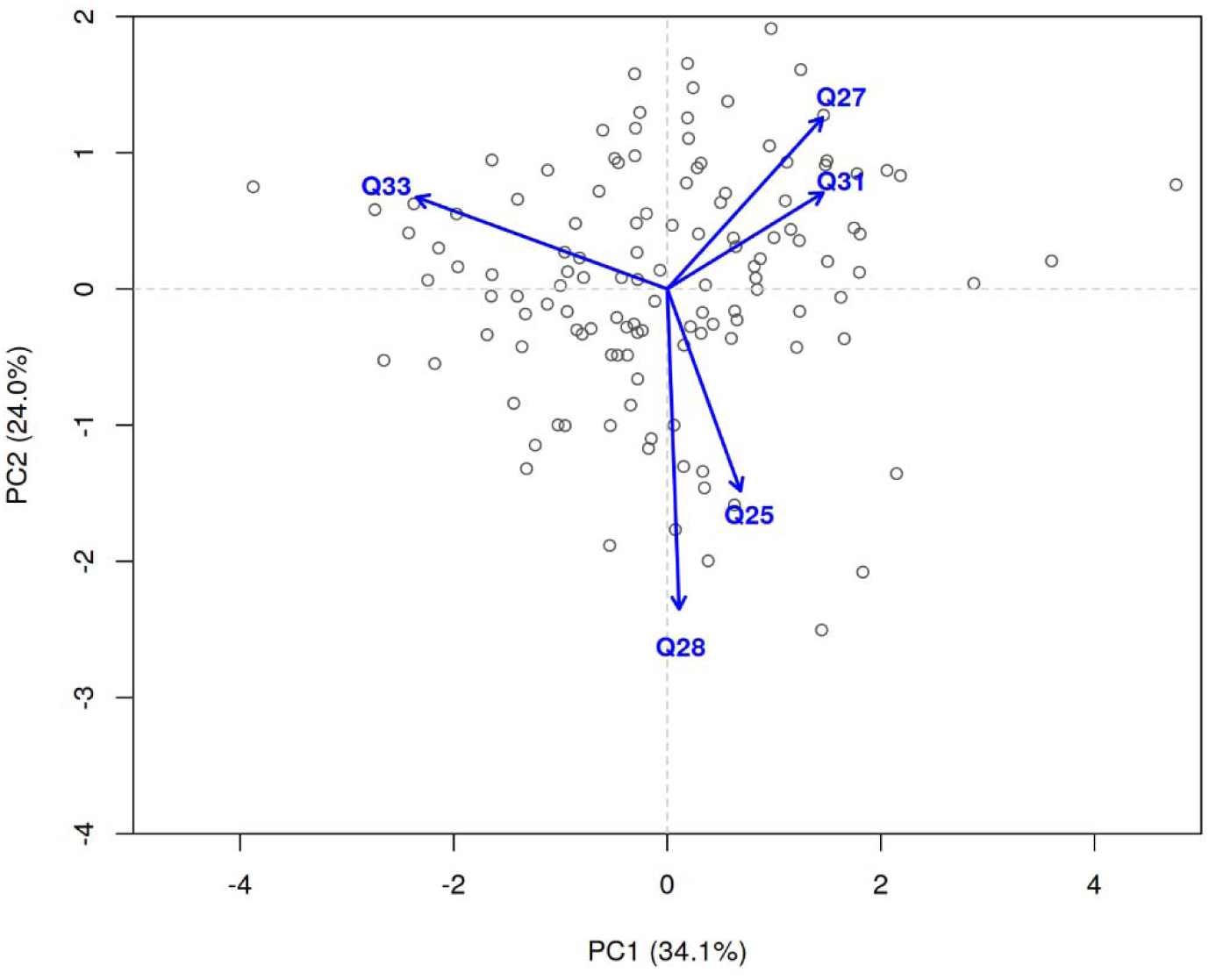
Principal component analysis (PCA) of *Clk* polyQ allele frequencies across European populations. Each black point represents a *Drosophila melanogaster* population, and blue arrows indicate the loadings of individual polyQ alleles. PC1 and PC2 together explain 58.1% of the total genetic variation in allele frequencies among populations. For clarity, one population with an extreme PC2 score (PC2 ≈ −7.8) falls below the plotted range; it was retained in all analyses.

### 3.2 Geographic and Environmental Correlates of *Clk* polyQ Allele Frequencies

To examine spatial patterns in Clk polyQ allele frequencies, we tested associations between allele frequencies and geographic variables (latitude, altitude and longitude). To separate spatial from seasonal variation, this analysis used only the 101 spring-collected populations. Because allele frequencies estimated from Pool-seq data are overdispersed relative to a binomial expectation (residual deviance approximately 2.0 to 4.6 times the residual degrees of freedom), the models were fitted with a quasibinomial error, which estimates the dispersion from the data and scales the standard errors accordingly. Among the five most common alleles, the longitudinal associations were the most robust: Q25 and Q27 were significantly associated with longitude (quasibinomial p = 0.0013 and p = 0.0044, respectively), Q25 becoming more frequent and Q27 less frequent towards the east (Figure 4). Q28 showed a weaker latitudinal association (p = 0.036), decreasing towards higher latitudes (Supplementary Figure S3). The remaining geographic and altitudinal effects, including those of Q31 and Q33, were not significant after accounting for overdispersion (all p > 0.12; Supplementary Figure S4) and are not interpreted further. These findings indicate spatially structured genetic variation across Europe along a predominantly east-west axis, consistent with potential clinal differentiation.

To interpret whether these spatial trends reflect underlying climatic differences, we summarized the main environmental gradients across Europe using principal component analysis of the bioclimatic dataset. The first two components (PC1 and PC2) together explained 73.3 percent of the total climatic variation (Supplementary Figure S5). PC1 represented a temperature–precipitation gradient, with high positive loadings for temperature variables and negative loadings for precipitation, such that higher PC1 values correspond to warmer and drier environments. PC2 captured a gradient of continentality, with strong positive loadings for minimum temperature of the coldest month, annual precipitation and a strong negative loading for temperature seasonality. High PC2 values describe regions with mild, wet winters and low seasonal temperature variation, while low PC2 values correspond to colder, more seasonal climates (Supplementary Table S2).

Geographic variables were closely correlated with these climatic axes (Supplementary Figure S6). As expected, PC1 decreased with latitude (t□□□ = –11.90, p < 2 × 10□¹□), confirming that southern populations experience warmer and drier conditions. More importantly, PC2 showed a pronounced negative association with longitude (t□□□ = –10.50, p < 2 × 10□¹□), revealing an east–west gradient in continentality. This longitudinal climatic gradient, which we previously highlighted as an underappreciated axis of environmental variation across Europe (Khatib et al., 2023, Kiat et al. 2021), provides an important spatial context for the variation observed in *Clk* allele frequencies.

Applying the same quasibinomial correction to the climatic models (Figure 3; Supplementary Table S3), the associations with PC2, the continentality axis, were the most robust. Q27 was positively associated with PC2 (OR = 1.228, quasibinomial p = 0.0004), indicating higher frequencies in colder, more seasonal eastern environments, whereas Q25 was negatively associated with PC2 (OR = 0.787, p = 0.0040), indicating higher frequencies in milder, western regions. The PC2 association of Q33 was not significant after correcting for overdispersion (OR = 0.916, p = 0.066) and is not emphasised.

**Figure 3.**
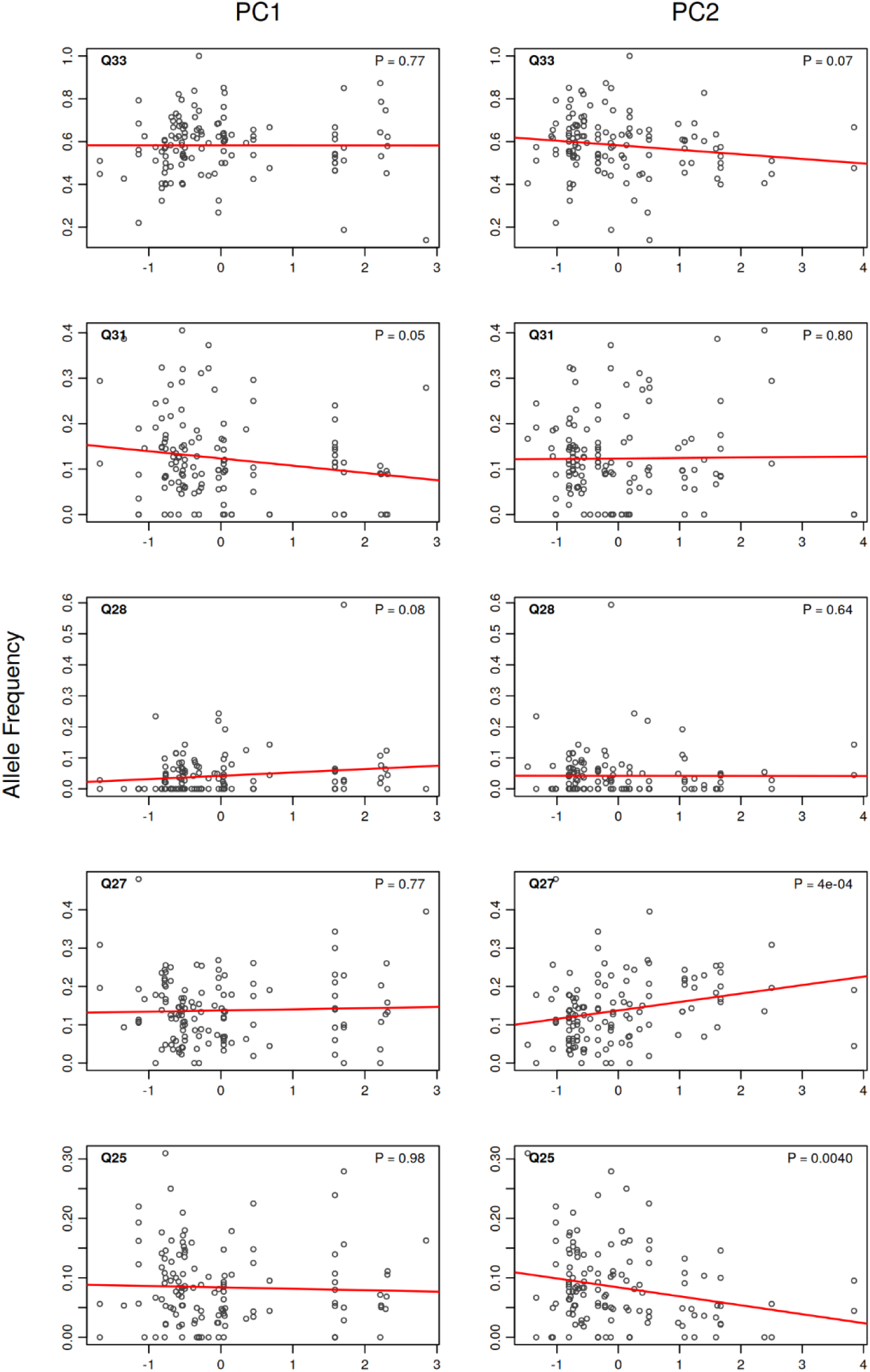
Spatial climatic gradients shape variation in CLK polyQ allele frequencies. PC1 captures a north–south gradient across Europe, while PC2 reflects an east–west gradient. Allele Q31 shows marginal significant associations with PC1, whereas Q25 and Q27 are significantly associated with PC2.

**Figure 4.**
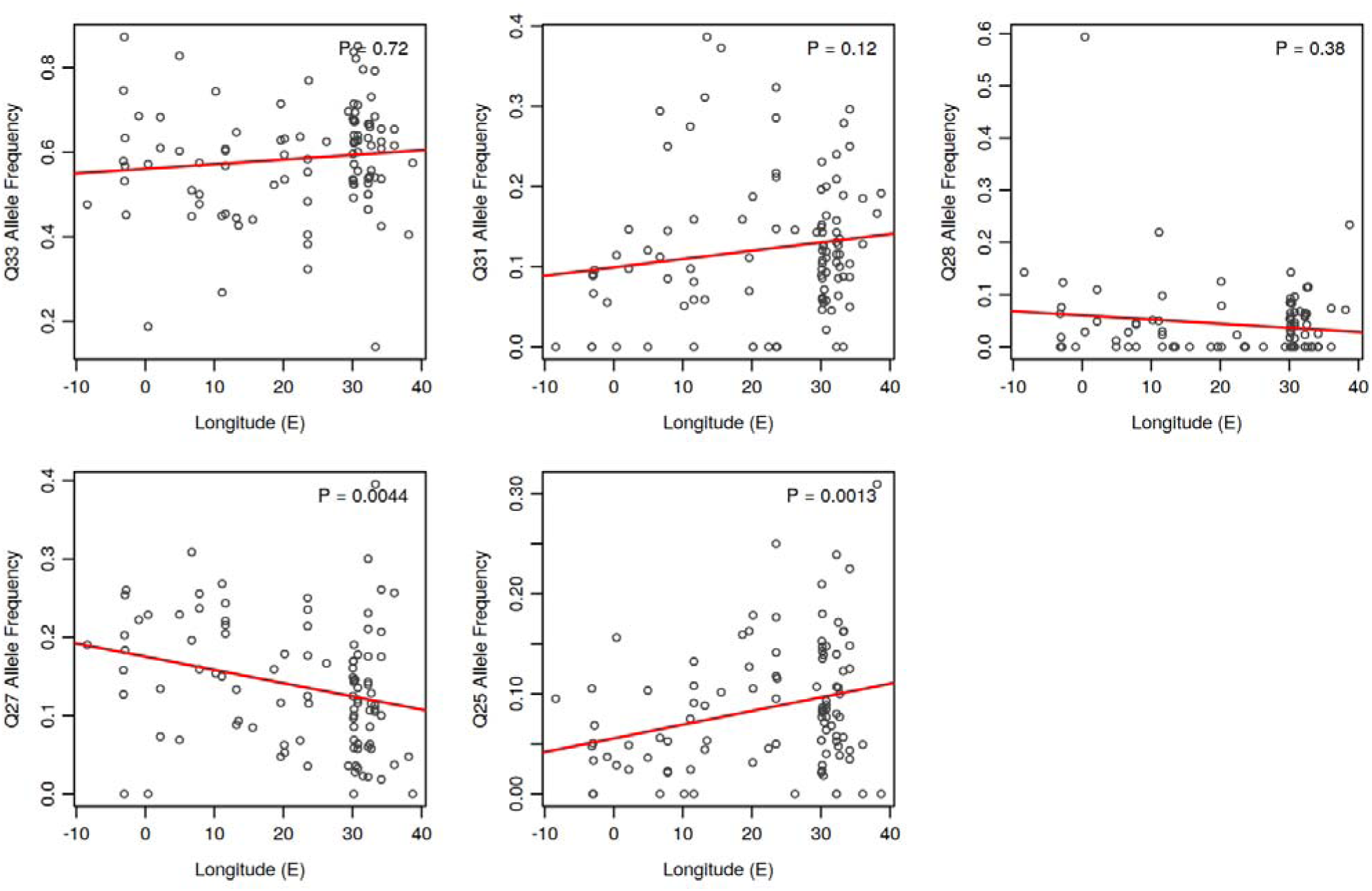
Correlations between polyQ alleles frequency with longitude across spring-collected populations (n = 101). Points are per-population allele frequencies; red lines are linear fits and P-values are from quasibinomial GLMs accounting for overdispersion. Q25 (P = 0.0013) and Q27 (P = 0.0044) are significant, defining the east–west cline.

The PC1 associations of Q31 and Q28 (OR = 0.862 and OR = 1.239 in the uncorrected binomial fit) did not remain significant after correcting for overdispersion (quasibinomial p = 0.050 and p = 0.080, respectively) and are therefore not interpreted as robust climatic clines.

Together, these analyses indicate that *Clk* polyQ allele frequencies exhibit spatial structuring across Europe, reflecting both geographic gradients and climatic variation summarized by PC1 and PC2. Although the effect sizes were modest, the consistent patterns across alleles suggest that temperature and seasonality may contribute to maintaining polymorphism at the *Clk* locus through spatially varying selection.

### 3.3 Analysis of Circadian Locomotor Activity

To evaluate whether variation in *Clk* polyQ length affects circadian temperature compensation, we measured locomotor activity rhythms in flies carrying different *Clk* polyQ alleles on a nearly isogenic background at 24 °C and 18 °C. After excluding individuals that died early, displayed arrhythmic activity, or showed extreme period values, 264 flies were included in the final analysis (12–36 individuals per allele per temperature; Figure5). Detailed mean ± SD values and sample sizes for each allele and temperature are provided in Supplementary Table S4. At 24 °C, flies carrying the Q21, Q25, and Q31 alleles exhibited slightly shorter mean free-running periods (FRPs) compared with 18 °C, whereas Q29, Q30, and Q33 showed little or no change.

A two-way ANOVA was performed with temperature, allele, and their interaction as fixed factors. There was no significant main effect of temperature on FRP (F□,□□□ = 2.42, p = 0.121), although a weak trend toward shorter FRP at higher temperature was observed (Figure 5). Neither the main effect of allele (F□,□□□ = 0.50, p = 0.779) nor the interaction between allele and temperature (F□,□□□ = 1.40, p = 0.224) was significant, indicating that FRP did not differ among polyQ variants and that temperature effects, where present, were consistent across alleles.

**Figure 5.**
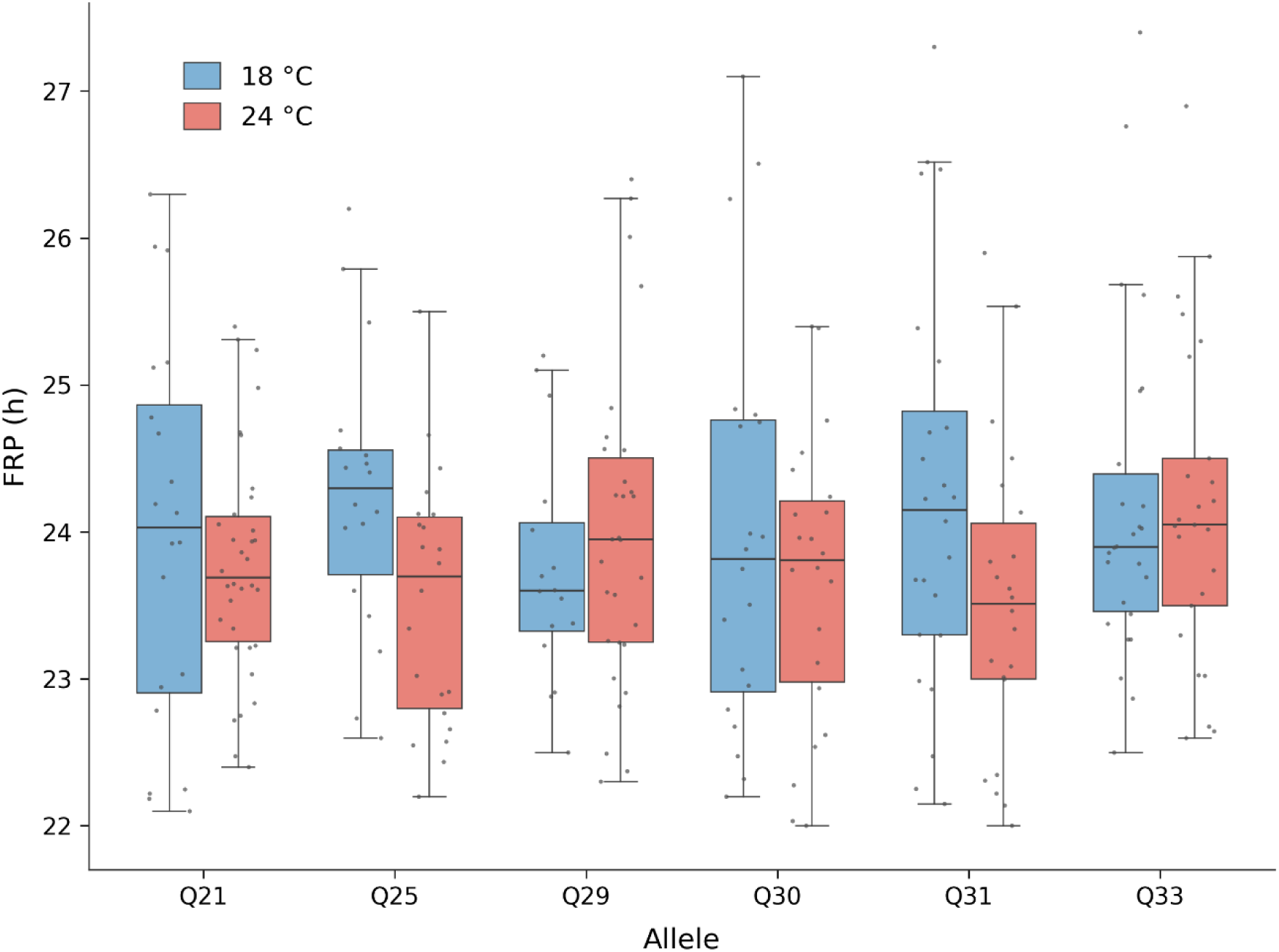
Effects of *Clk* polyQ variation and temperature on circadian free-running period (FRP). Boxplots show the distribution of FRP (hours) for flies carrying six *Clk* polyQ alleles (Q21, Q25, Q29, Q30, Q31, Q33) measured at 18 °C (blue) and 24 °C (red) under constant darkness. Each point represents an individual fly. Boxes indicate the interquartile range, horizontal lines mark medians, and whiskers extend to 1.5 × the interquartile range. The plot illustrates variation in circadian period across genotypes and temperatures. Detailed sample sizes and descriptive statistics are provided in Supplementary Table S4.

To assess temperature compensation at the level of individual genotypes, we next examined within allele changes in FRP using a Bayesian model (Table 1). For each allele, the magnitude of temperature dependent change in FRP (ΔFRP, 18 °C to 24 °C) was summarized by the posterior mean and 95 percent credible interval (CI). Q25 showed a credible shortening of FRP at higher temperature (posterior mean ΔFRP = −0.68 h, 95% CI = −1.27 to −0.09, P(Δ < 0) = 0.99). Q31 exhibited a moderate decrease in FRP, although with greater uncertainty (ΔFRP = −0.52 h, 95% CI = −1.16 to 0.11, P(Δ < 0) = 0.95), with a similar but weaker pattern for Q30 (ΔFRP = −0.42 h, 95% CI = −1.18 to 0.34, P(Δ < 0) = 0.86). For the remaining alleles, credible intervals overlapped zero, indicating no detectable temperature dependent change in FRP. Pairwise comparisons supported a stronger temperature response in Q25 relative to most other alleles, although overlap among posterior distributions indicated that differences were modest overall (Supplementary Table S5).

**Table 1.** Bayesian model estimates for ΔFRP (18 °C → 24 °C) across *Clk* polyQ alleles. Posterior mean ΔFRP, 95% credible interval (CI), and posterior probability P(Δ < 0).

| Allele | ΔFRP (h) | 95% CI (h) | $P(\Delta < 0)$ |
| --- | --- | --- | --- |
| Q21 | -0.14 | [-0.71, 0.43] | 0.68 |
| Q25 | -0.68 | [-1.27, -0.09] | 0.99 |
| Q29 | 0.31 | [-0.25, 0.87] | 0.14 |
| Q30 | -0.42 | [-1.18, 0.34] | 0.86 |
| Q31 | -0.52 | [-1.16, 0.11] | 0.95 |
| Q33 | 0.23 | [-0.46, 0.92] | 0.25 |

Overall, these results indicate that FRP was only weakly influenced by temperature across alleles. Among the variants examined, Q25 showed the strongest evidence for temperature sensitivity, consistent with a modest reduction in temperature compensation relative to the other alleles.

We next examined whether polyQ length affects the timing of locomotor activity under LD. The day-folded average activity profiles are presented in Supplementary Figure S8. Although phase differences are modest in magnitude, statistical analysis using seven pre-specified phase references revealed significant differences (Supplementary Table S6). Temperature had a strong and consistent effect on every phase reference (two-way ANOVA, main effect of temperature P < 10□□ for all seven references; e.g. evening peak F_1,736_ = 166.6, P = 1.6 × 10□³□). Cooling from 24 °C to 18 °C advanced the evening peak (from a genotype range of ZT 11.3 to 12.0 down to ZT 10.9 to 11.1) and delayed the morning peak (from ZT 0.4 to 1.1 up to ZT 1.2 to 1.5), so that the two activity components moved closer together and the morning-to-evening interval contracted (from ZT 10.3 to 11.6 at 24 °C to ZT 9.5 to 9.8 at 18 °C). These shifts were common to all genotypes.

Against this background, polyQ genotype affected activity phase specifically at the warmer temperature. At 24 °C the genotypes differed significantly in all three evening references and in the morning-to-evening interval (one-way ANOVA: evening onset F_5,346_ = 7.25, P = 1.8 × 10□□; evening peak F_5,346_ = 9.48, P = 1.7 × 10□□; evening offset F5,346 = 4.33, P = 7.7 × 10□□; morning-to-evening interval F_5,346_ = 8.87, P = 6.1 × 10□□; morning peak F_5,346_ = 5.98, P = 2.5 × 10□□; all significant after Benjamini-Hochberg correction across the seven references; Supplementary Table S6). Post hoc comparisons (Tukey HSD) resolved two consistent groups: Q29 and Q33, together with Q25, showed the latest evening peaks and the longest morning-to-evening intervals, whereas Q21 and Q30 showed the earliest evening peaks and the shortest intervals, with Q31 intermediate (Figure 5; Supplementary Table S6). The effects were modest in magnitude, with genotype explaining approximately 10 to 12 % of the among-fly variance in the strongest references (evening peak and interval) and with substantial overlap among individuals; consequently these differences were not apparent in the population-average waveforms and emerged only from the per-fly analysis. At 18 °C, by contrast, no phase reference differed among genotypes after correction, and the genotype means converged onto a common phenotype (Figure 5; Supplementary Table S6).

This temperature dependence was confirmed by a significant genotype by temperature interaction, tested with a distribution-free Freedman-Lane permutation procedure. The interaction was significant for the evening references and the morning-to-evening interval (evening peak and interval P = 0.002, evening onset P = 0.006, evening offset P = 0.037, Benjamini-Hochberg corrected) and marginal for the morning peak (P = 0.050), but absent for the morning onset and offset (Supplementary Table S6). Variation in *Clk* polyQ length therefore has little effect on the phase of activity under cooler conditions but produces reproducible differences in the timing of the evening component under warmer conditions, primarily by advancing or delaying the evening peak and thereby narrowing or widening the interval between the morning and evening activity bouts.

Overall, these analyses indicate that variation in *Clk* polyQ length does not strongly alter temperature compensation of circadian period, as FRP remained largely stable across alleles. In contrast, circadian phase exhibited greater sensitivity to temperature, with pronounced allele specific differences emerging at elevated temperature. Among the variants examined, Q25 showed the clearest reduction in temperature compensation of period, whereas the temperature-dependent divergence in phase was greatest for Q29 and Q33, which displayed the latest evening activity and the longest morning-to-evening phase angle at 24 °C.

### 3.4 Effects of *Clk* polyQ Variation on Transcriptional Activity

Luciferase reporter assays were used to assess how variation in *Clk* polyQ length influences CLK transcriptional activity. Control transfections containing the empty expression vector showed very low relative luciferase activity, approximately 95-fold lower than that observed for the least active CLK variant. The analysis revealed a significant overall effect of polyQ length on reporter expression (Kruskal–Wallis test: χ² = 14.58, df = 3, p = 0.002; Figure 6). Pairwise comparisons indicated that the longer alleles, polyQ31 and polyQ33, exhibited significantly higher transcriptional activity than polyQ21 (p = 0.015 for both) and polyQ28 (p = 0.015 and p = 0.023, respectively). No significant differences were found between polyQ31 and polyQ33 or between polyQ21 and polyQ28. These results show that CLK variants with longer polyQ tracts promote stronger activation of the *per* E-box reporter, suggesting that natural variation in polyQ length modulates CLK’s transcriptional potency.

**Figure 6.**
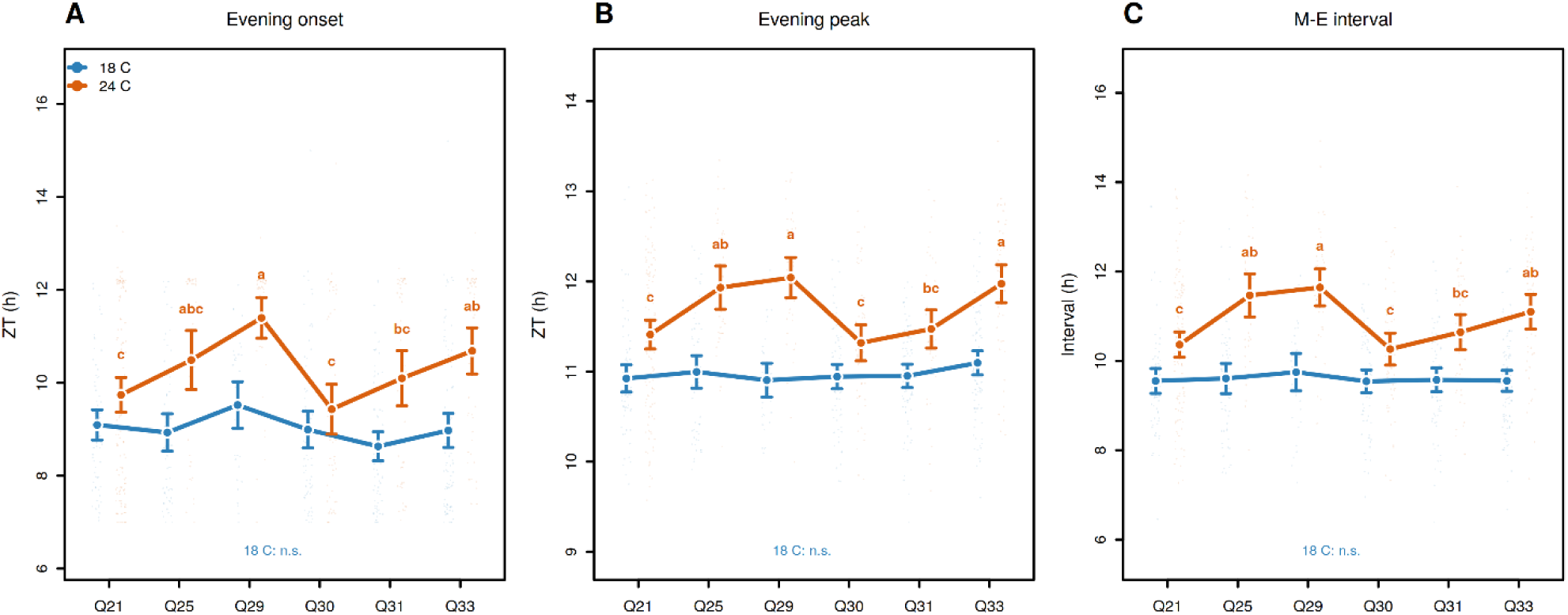
*Clk* polyQ genotype shapes evening phase timing at 24 °C but not at 18 °C. Evening onset (A), evening peak (B), and the morning to evening interval (C), expressed in Zeitgeber time (ZT, hours), for six *Clk* polyQ genotypes (Q21, Q25, Q29, Q30, Q31, Q33) recorded under 12:12 LD at 18 °C (blue) and 24 °C (orange). Faint points are individual flies; filled symbols show genotype means with 95% confidence intervals (n = 352 flies at 24 °C, 396 at 18 °C). At 24 °C, genotype means sharing a letter do not differ (Tukey HSD, P < 0.05); at 18 °C the genotypes are statistically indistinguishable (n.s.). The divergence of genotypes at 24 °C together with their convergence at 18 °C reflects a significant genotype by temperature interaction for all three references (Freedman-Lane permutation, P < 0.05).

**Figure 7.**
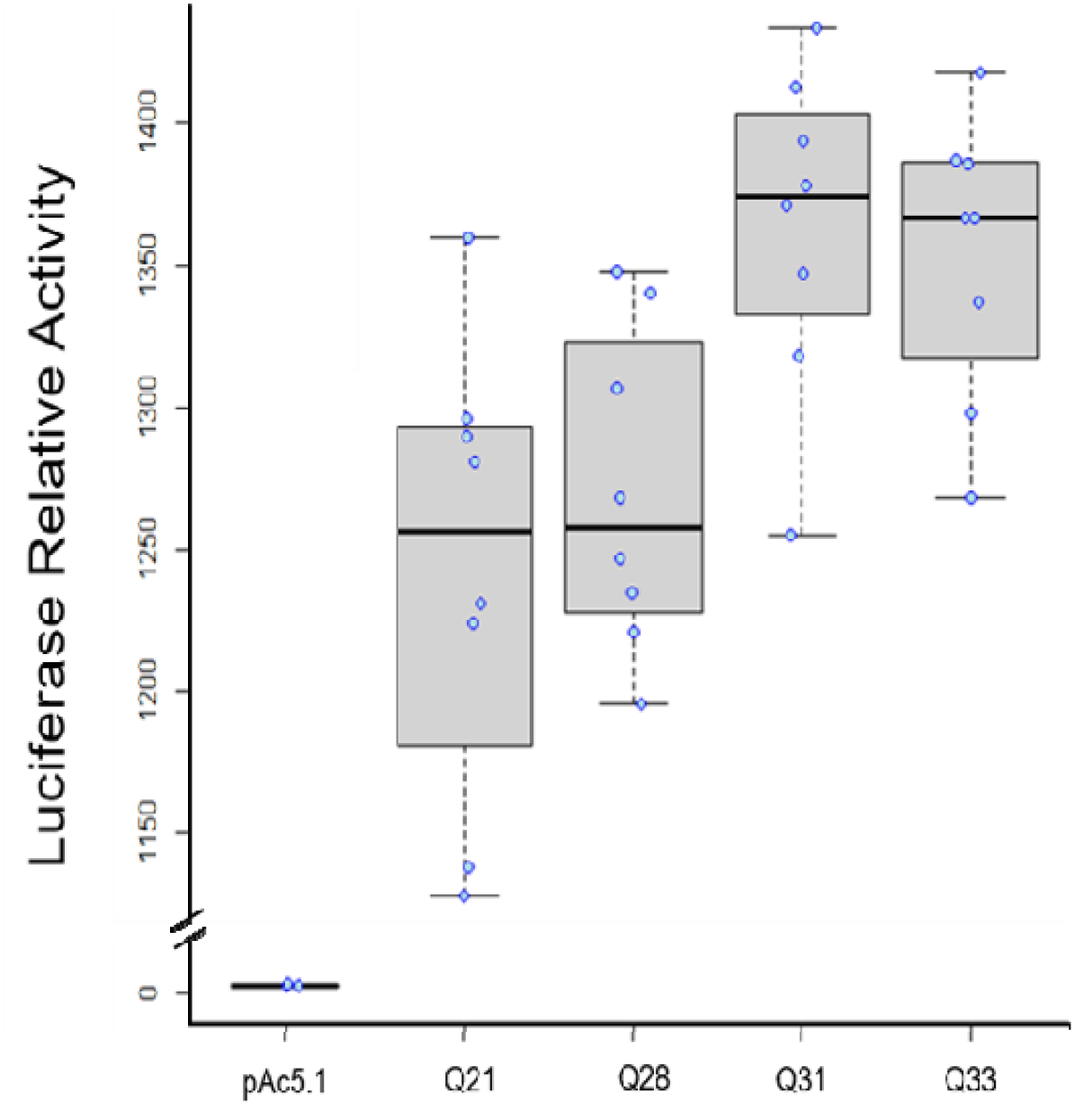
Variation in CLK polyQ length influences transcriptional activity in a luciferase reporter assay. Luciferase activity was measured for different CLK polyQ alleles (Q21, Q28, Q31, Q33) and the pAc control. Each point represents an independent biological replicate (eight for each allele, three for the control). PolyQ31 and PolyQ33 showed significantly higher activity than PolyQ21 and PolyQ28 (p < 0.05, Wilcoxon test with Benjamini–Hochberg correction), while no significant differences were detected between PolyQ31 and PolyQ33 or between PolyQ21 and PolyQ28.

Overall, these results indicate that CLK variants with longer polyQ tracts exhibit enhanced transcriptional activation of the *per* E-box reporter, consistent with length-dependent modulation of CLK activity.

## **4** Discussion

Natural variation in circadian clock genes provides a valuable framework for understanding how organisms adapt to environmental heterogeneity. In *Drosophila melanogaster*, polymorphisms in the *Clock (Clk)* gene are of particular interest since CLK plays a central role in the transcriptional feedback network that sustains circadian oscillations. This study reveals that variation in the length of the polyQ tract within *Clk* contributes to both the behavioral and molecular tuning of the circadian system. Using European population-genomic data, behavioral assays in near-isogenic lines, and luciferase reporter analyses, we found that *Clk* polyQ alleles differ in frequency across climatic gradients and exhibit functional variation under laboratory conditions. These findings reveal potential mechanisms through which environmental variation could influence allele maintenance in natural populations.

Patterns of *Clk* allele distribution across Europe structured along geographic axes. The most robust clines were longitudinal: Q27 was more frequent in eastern, more seasonal regions and Q25 in milder, western regions, whereas the weaker latitudinal and altitudinal trends of the longer alleles Q31 and Q33 did not remain significant after accounting for overdispersion. Although longitude is not itself a selective agent, in Europe it corresponds to a gradient of continentality, from the mild oceanic west to the colder, more seasonal east, captured here by PC2 (Supplementary Table S2). We have previously documented longitudinal clines in circadian clock genes along this frequently overlooked axis (Khatib et al., 2023), so a longitudinal cline in *Clk* polyQ frequency is consistent with a real environmental gradient, though it could also partly reflect colonisation history. The longitudinal cline in Q25 and Q27 aligns with this environmental axis; however, such spatial patterns can arise from colonisation history, demographic processes, or selection, and our data do not permit distinguishing among these mechanisms. More broadly, clinal variation in clock genes has been reported across taxa, including the *per* Thr–Gly polymorphism (Costa et al., 1992) and *Clk* polyQ variation in salmon (O’Malley & Banks, 2008) and birds (Johnsen et al., 2007). Although the associations between allele frequencies and climatic or geographic gradients were generally weak to moderate, their consistency across datasets and correspondence with environmental axes warrant further investigation into potential selective mechanisms, possibly linked to the thermal and photoperiodic stability of local environments.

The polyQ polymorphism in *Drosophila melanogaster* was previously examined in Australian populations, where Weeks et al. (2006) reported no evidence of a cline in *Clk* repeat length along the east coast. In the same study, they also failed to detect a latitudinal pattern in the *period (per)* Thr–Gly repeat, a result that contrasted with the strong European cline originally described by Costa et al. (1992). This absence of a pattern in Australia prompted debate about whether clinal variation in clock genes truly reflects adaptive differentiation or arises from sampling and methodological artefacts. However, subsequent analyses by Kyriacou et al. (2007) and Sawyer et al. (2006) re-examined Australian and African data and identified a significant, though weaker, cline in *per* that was consistent in direction with the European gradient, supporting the thermal-adaptation hypothesis. These authors argued that the apparent lack of a cline in Weeks et al. (2006) likely resulted from limited allele resolution, potential species misidentification, and the relatively recent colonization history of *D. melanogaster* in Australia, rather than the absence of selection. This historical debate over *per* underscores how demographic history and sampling scale can obscure adaptive spatial patterns, a consideration directly relevant to interpreting the geographic structuring of *Clk* polyQ variation in our European dataset.

Behavioral analyses of near-isogenic lines showed that most *Clk* polyQ variants maintained a largely stable free-running period (FRP) across 18 °C and 24 °C, indicating effective temperature compensation. However, flies carrying the Q25 allele displayed a consistent shortening of FRP at higher temperature, suggesting reduced compensation. This pattern is analogous to that observed for *per* Thr–Gly polymorphisms, where specific repeat lengths altered thermal robustness (Sawyer et al., 1997). Interestingly, both the shortest (Q21) and longest (Q33) alleles exhibited strong compensation, while intermediate alleles, particularly Q25, were more temperature-sensitive, revealing a non-linear relationship between polyQ length and circadian stability. In contrast to these allele specific effects on circadian traits under laboratory conditions, circadian phase did not show a consistent relationship with geographic variables, suggesting that selection on *Clk* polyQ length is unlikely to act directly on phase timing alone.

At the molecular level, luciferase reporter assays demonstrated that CLK transcriptional activity also depends on polyQ length. Longer variants (Q31, Q33) exhibited higher activation of a *per* E-box reporter compared with shorter alleles (Q21, Q28), suggesting that polyQ extension enhances CLK’s ability to promote target gene expression. Similar effects of polyQ length on transcriptional activation have been reported in other regulatory proteins (Atanesyan et al., 2012; Gemayel et al., 2015). The correspondence between transcriptional strength and aspects of behavioral robustness suggests that differences in CLK activity, specifically the higher transactivation of longer polyQ variants, could contribute to circadian function. Testing CLK transactivation under varying thermal conditions would be necessary to directly assess whether transcriptional differences mediate temperature compensation. The transcriptional constructs (Q21, Q28, Q31 and Q33) were selected a priori from allele frequency and PCA clustering, before the behavioural assays identified Q25 as the temperature-sensitive allele; a Q25 construct is therefore a priority for follow-up work. Notably, although the Q21 variant carries an internal histidine that interrupts its glutamine tract, its transcriptional activity was comparable to that of the other short allele, Q28, consistent with the length of the uninterrupted glutamine stretch, rather than the total repeat region, being the primary determinant of transcriptional activity.

With respect to circadian phase specifically, the transcriptional and behavioral phenotypes are best viewed as complementary rather than directly predictive. Among the alleles common to both assays, reporter activity and evening phase did not co-vary monotonically, indicating that steady state transactivation alone does not set phase. A plausible link is that differences in CLK transactivation alter the kinetics of the PER and TIM feedback loop, and thus the timing of the evening peak, so that reporter magnitude and phase shift need not be tightly coupled; time resolved assays capturing oscillation phase will be needed to test this.

The physical basis of these functional effects likely involves the intrinsic structural properties of polyQ domains. PolyQ sequences facilitate multivalent, weak interactions that promote the formation of dynamic, membraneless condensates through liquid–liquid phase separation (Banani et al., 2017; Peng et al., 2020; Kawasaki & Fukaya, 2023). In the *Arabidopsis* clock protein ELF3, phase separation of the prion-like domain is driven by rising temperature, so that warming, rather than cooling, promotes condensate formation, and lengthening the polyQ tract broadens the range of temperature and pH over which the condensed phase forms and persists while having a comparatively small effect on the transition temperature itself (Hutin et al., 2023). By analogy, longer *Clk* polyQ alleles may promote and stabilize CLK condensates across a broader range of thermal conditions and thereby support stronger transactivation, consistent with the higher reporter activity we observed for Q31 and Q33, whereas shorter tracts may sustain condensates over a narrower range. This interpretation is consistent with biophysical studies showing that shorter polyQ tracts undergo greater temperature-induced conformational changes (Cui et al., 2014).

This mechanism appears to be a broadly conserved feature of circadian clock architecture: a recent survey of intrinsically disordered regions across fungal, insect, and mammalian core clock proteins highlights how such disordered, low-complexity domains, including polyQ and prion-like tracts, function as tunable, temperature-sensitive interaction modules that support condensate assembly and modulate clock protein activity (Usher & Pelham, 2026).

The unexpected abundance of the temperature-sensitive Q25 allele in eastern populations may reflect additional or indirect selective pressures. One possibility is that greater structural flexibility of Q25 confers advantages for other temperature-linked traits such as diapause or cold response, both regulated by circadian pathways (Hidalgo & Chiu, 2024; Goto et al., 2006; Cai et al., 2024). Alternatively, heterozygosity for Q25 and longer alleles like Q33 could produce composite phenotypes with broader environmental tolerance, maintaining diversity through balancing selection. Similar maintenance of *Clk* polyQ diversity has been reported in migratory birds exposed to variable photoperiods (Justen et al., 2022).

Beyond repeat length, additional molecular features likely shape CLK function. The polyQ region can be encoded by mixed CAG and CAA codons, whose differing translation speeds influence protein folding and stability (Fu et al., 2016; Moreno-Rodríguez et al., 2025). Moreover, the structure of neighboring domains modulates the conformational flexibility and accessibility of the polyQ tract (Mier & Andrade-Navarro, 2021). Such interactions may explain why alleles of similar length, such as Q21 and Q33, exhibit comparable temperature robustness, whereas Q25 shows distinct behavior. Detailed analyses of codon composition, adjacent sequence variation, and post-translational modifications will be essential to fully understand how natural *Clk* variants influence clock performance.

In summary, our results demonstrate that *Clk* polyQ length polymorphism affects both the molecular activity of CLK and the thermal stability of circadian behavior in *Drosophila melanogaster*. The geographic distribution of alleles, particularly the pronounced longitudinal cline in Q25 and Q27, is structured in patterns consistent with underlying environmental variation, though demographic and selective mechanisms cannot be distinguished with current data. The non-linear relationship between polyQ length and function highlights the adaptive potential of low-complexity regions, which can modulate protein dynamics and environmental responsiveness without major changes in overall structure. These findings contribute to a broader understanding of how subtle molecular variation in key regulatory proteins underpins the evolution of environmental adaptation in circadian systems.

## **6** Author Contributions

M.Y. conducted the experiments, analyzed the data, and wrote the first draft of the manuscript. B.F. assisted with the experiments and supervised the experimental work. M.K. assembled and processed the sequencing data. E.T. designed the research and contributed to data analysis. All authors revised the manuscript and approved the final version.

## **8** Ethics Statement

All experiments in this study were performed on the invertebrate *Drosophila melanogaster*, an organism that is not subject to the ethical regulations governing vertebrate animal research. No ethical approval or license was therefore required for this work.

## Data Availability Statement

All data, analysis scripts (R and RMarkdown), and intermediate datasets are archived at Zenodo (https://doi.org/10.5281/zenodo.22030098).

