## Supplementary Figure for "Natural Variation in *Drosophila melanogaster Clock* PolyQ Length: Geographic Gradients and Functional Properties"

**Supplementary Material**

Natural Variation in *Clock* polyQ Length Is Associated with Circadian Function to Climatic Gradients in *Drosophila melanogaster*

Maya Yair^1^, Bettina Fishman^1^, Martin Kapun^2^ and Eran Tauber^1*^

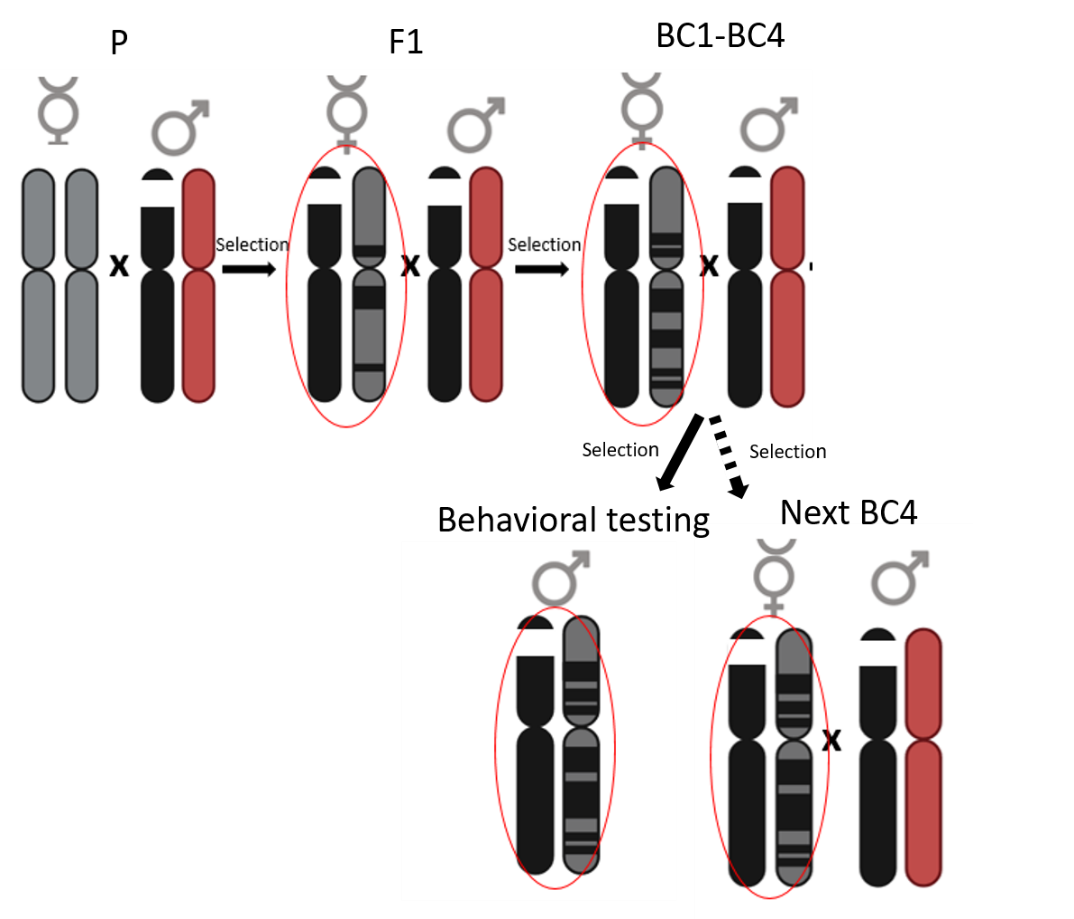

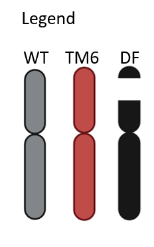

Supplementary Figure S1. Crossing scheme for generating the polyQ NILs. The 3rd chromosome consists of WT 3L (grey), TM6C balancer (red), and the chromosomes with the deficiency (Df(3L)pbl-X1; black(. An initial cross between deficiency- carrying males, and WT females from each polyQ line was conducted. Hemizygote virgin females with WT bristles (no balancer) were collected from F1 offspring and backcrossed with w [1118]; Df(3L)pbl-X1/TM6C males to generate BC1. The following BC2-BC4 were generated by backcrossing white-eyed hemizygote virgin females offsprings of the previous backcross with the deficiency-carrying males three more times. In each step, the contribution of the donor parent genome is reduced by half, so that BC4 flies have 96% genomic similarity. From each BC4 population, hemizygote white-eyed and WT bristled females and males were collected. Hemizygous males were tested for their circadian activity while the females were crossed to generate the next BC.

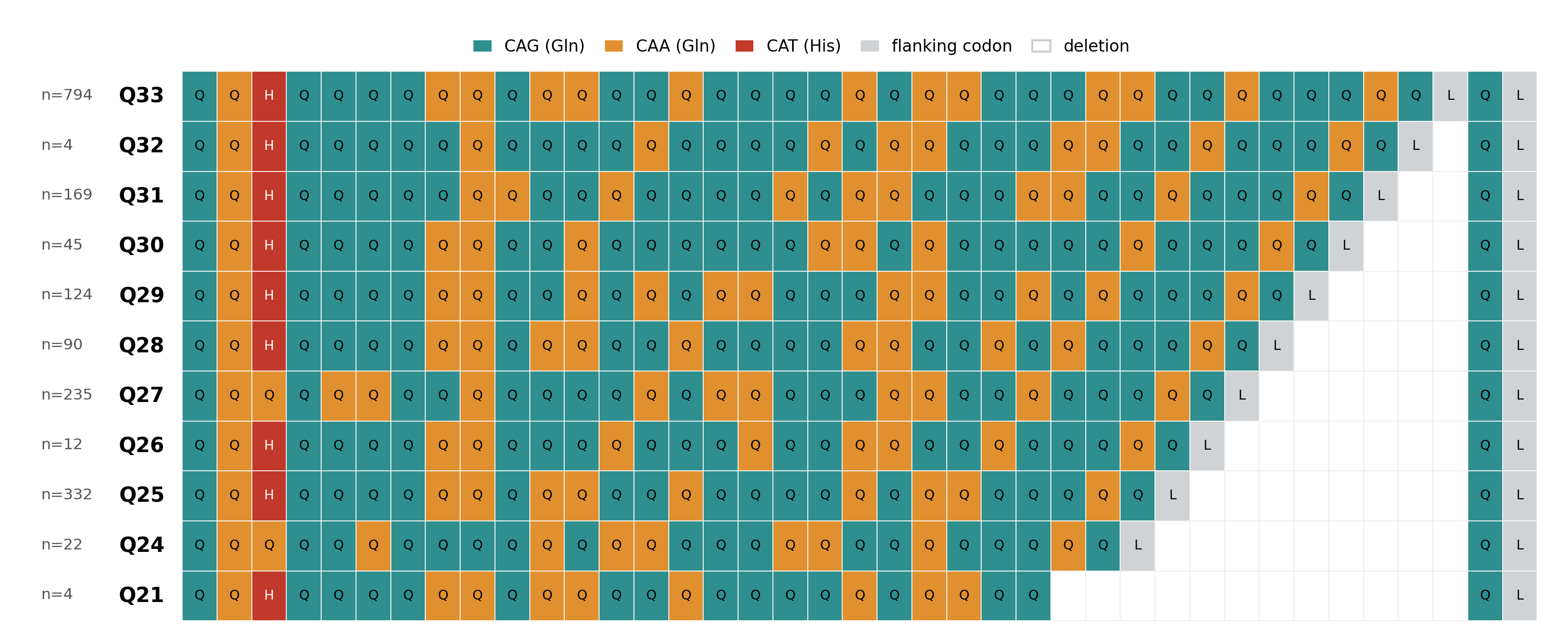

****Supplementary Figure S2. Sequence variation among the eleven**** Clk ****polyQ length alleles in European**** Drosophila melanogaster****.**** Coding-strand codon alignment (N to C terminus) of the eleven length variants (Q21 to Q33) recovered from DrosEU Pool-seq data. Each cell represents one codon, coloured by identity: CAG (teal) and CAA (amber) both encode glutamine, showing the synonymous codon interspersion within the tract; CAT (red) encodes a histidine near the tract margin, present in nine alleles and replaced by glutamine in Q24 and Q27; flanking codons are shown in grey. Length differences relative to the longest allele (Q33) are drawn as a single nested deletion block; because deletions cannot be localised within a tandem repeat, gap placement within the tract is arbitrary. The number of spanning reads supporting each allele is given at left; Q21 and Q32 (n = 4) are the least supported.

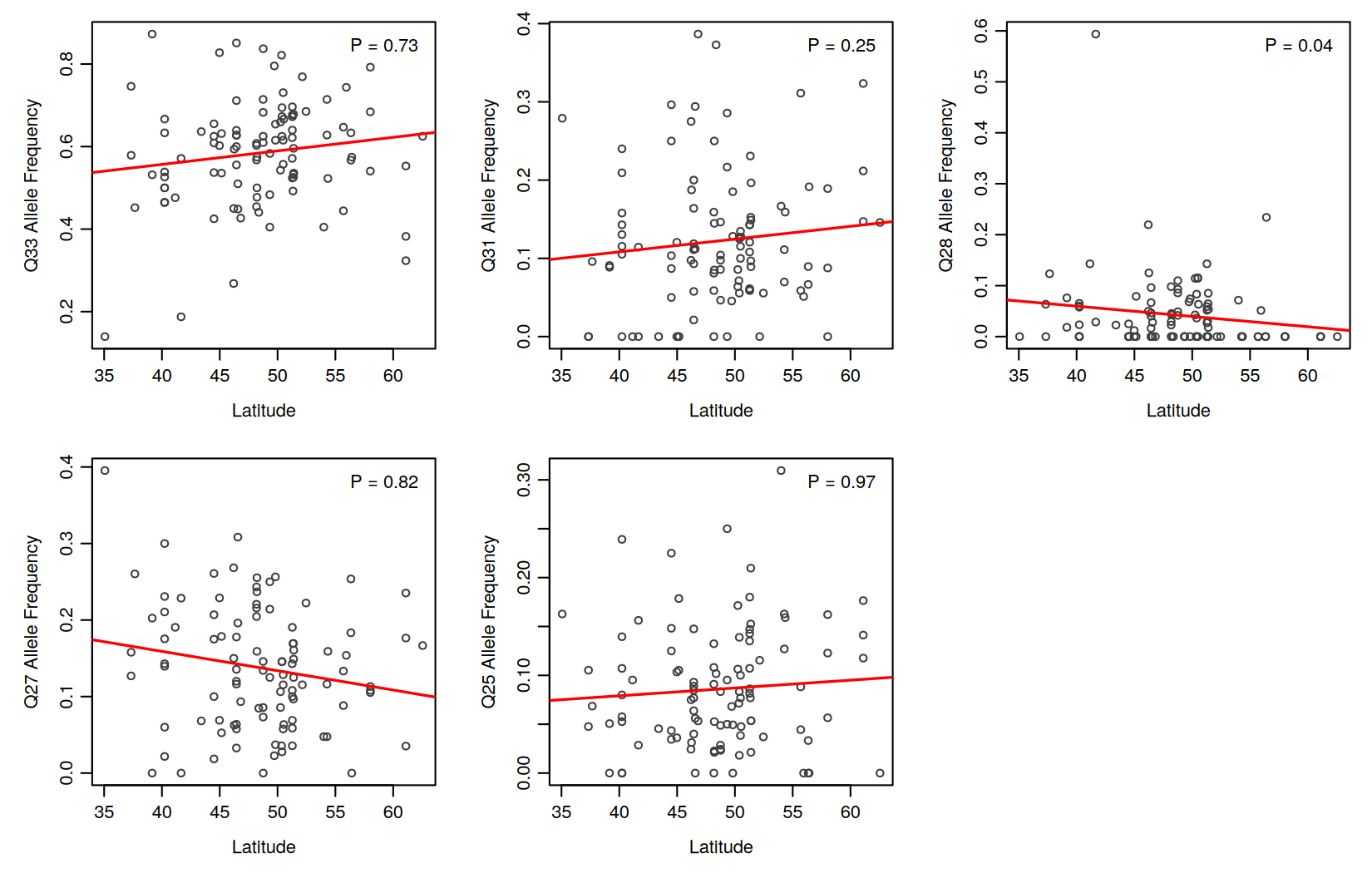
Supplementary Figure S3. Correlations between *Clk* polyQ allele frequencies and latitude across spring-collected populations (n = 101). Points are per-population allele frequencies; red lines are linear fits and P-values are from quasibinomial GLMs accounting for overdispersion. Only Q28 is significant (P = 0.036).

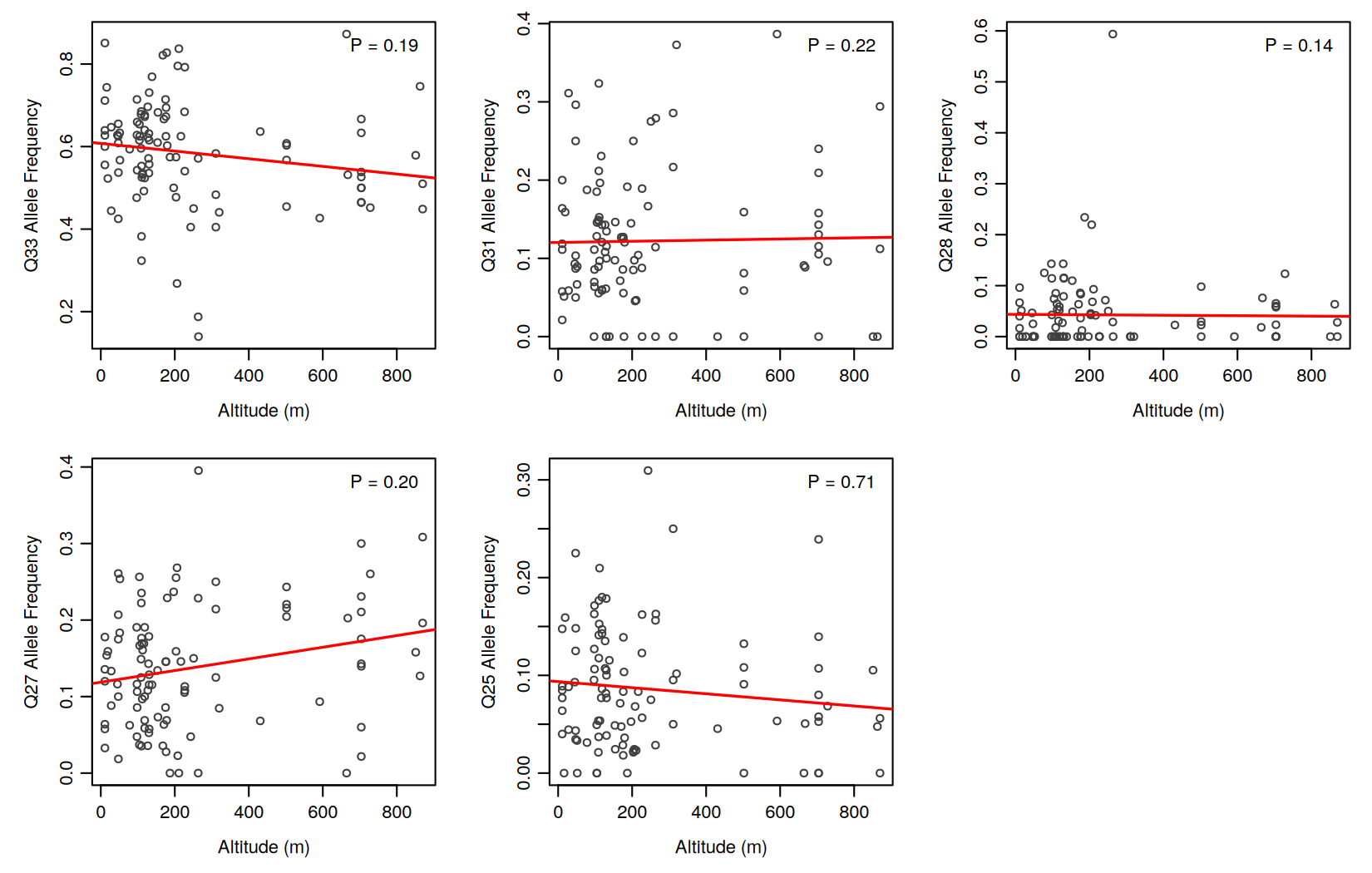

Supplementary Figure S4. Correlations between polyQ alleles frequency with altitude across spring-collected populations (n = 101). Points are per-population allele frequencies; red lines are linear fits and P-values are from quasibinomial GLMs accounting for overdispersion. No allele is significant after correction.

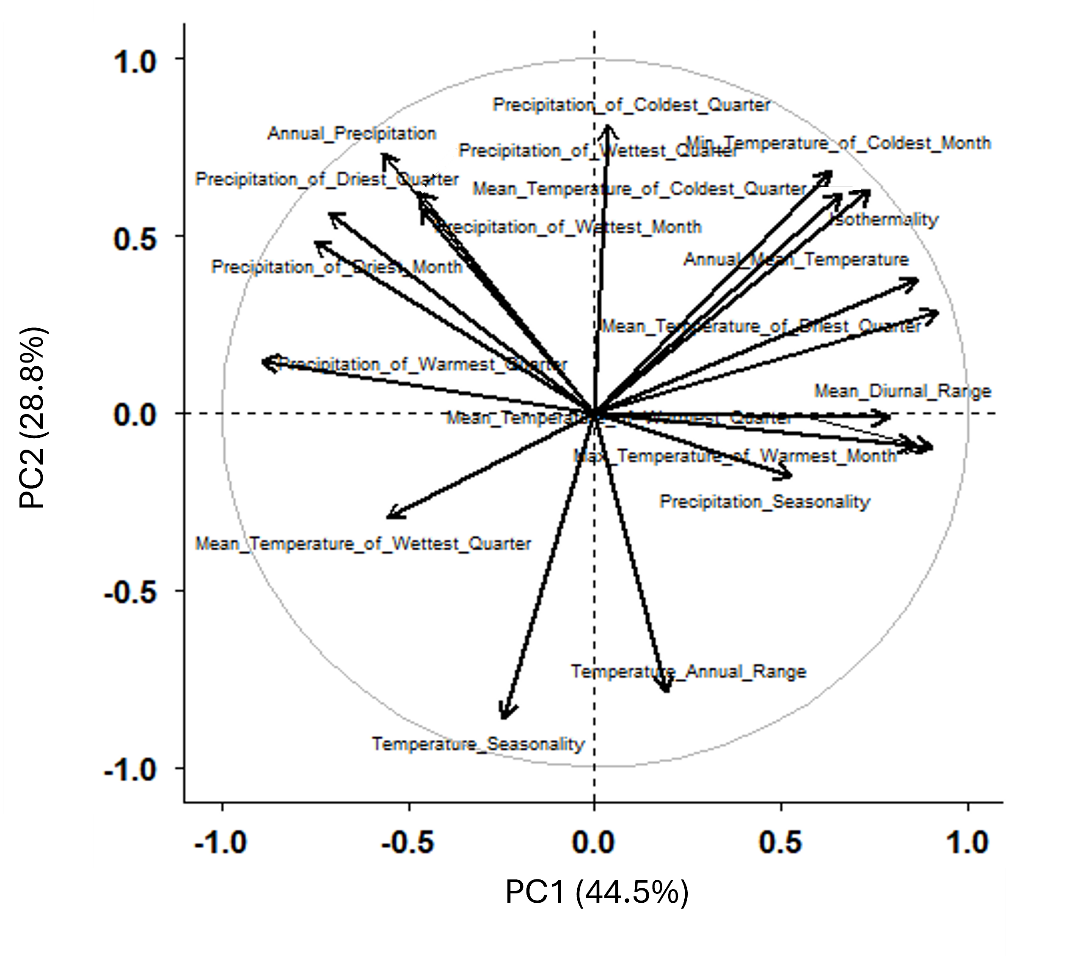

Supplementary Figure S5. PCA analysis of 19 Bioclimatic variables measured across Europe, corresponds to locations of DrosEU *drosophila* populations. The loading of each of the variables is shown by black arrow. PC1 and PC2 explain 73.3 % of the variation across the population.

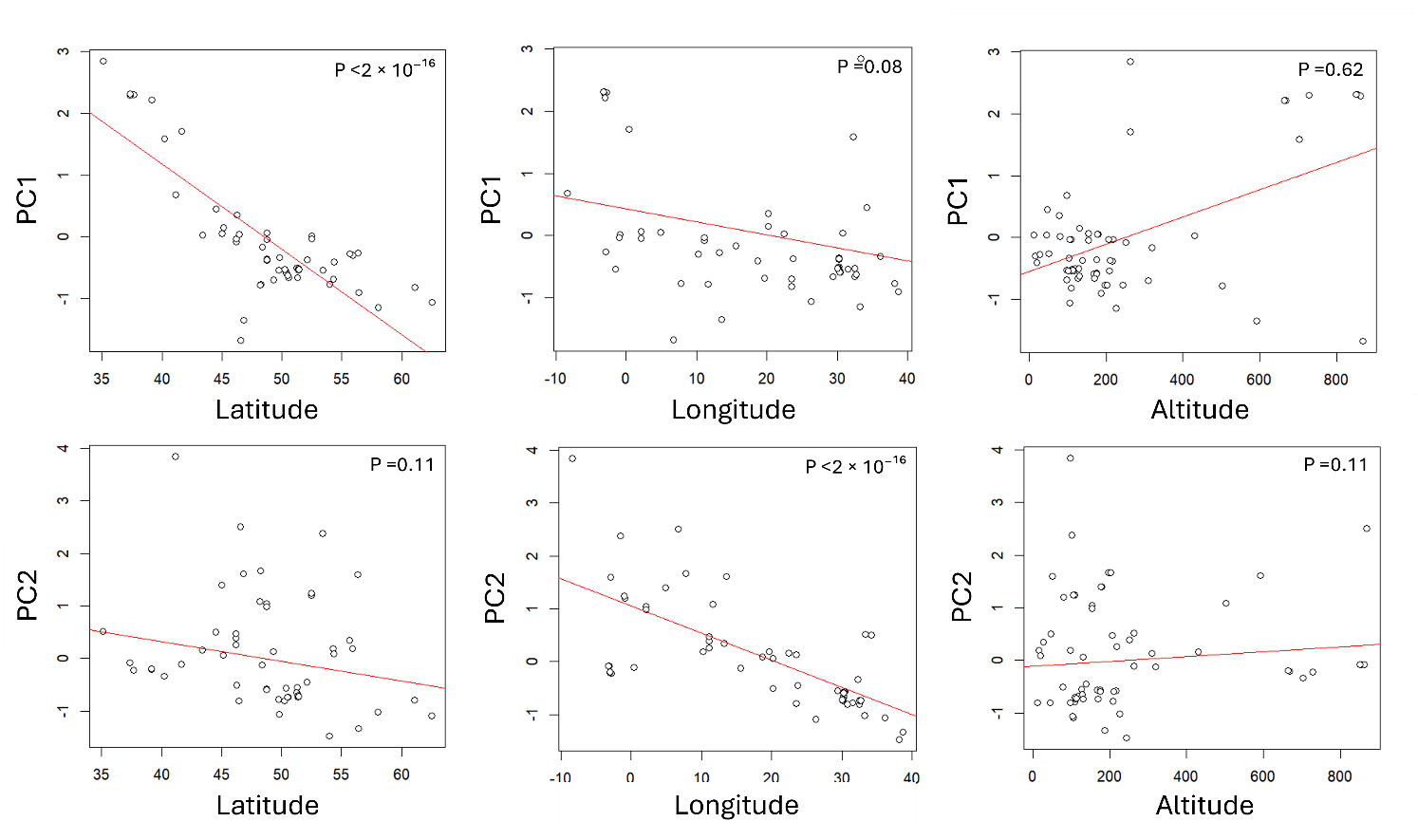

Supplementary Figure S6. Principal components (PC1 and PC2) of 19 bioclimatic variables and their correlations with latitude, longitude, and altitude. PC1, strongly correlated with latitude, represents a north–south gradient in temperature and precipitation, with high values (south) indicating warmer, drier climates and low values (north) cooler, wetter ones. PC2, correlated with longitude, reflects an east–west gradient from mild, humid western regions to colder, more seasonal eastern environments.

****Supplementary Figure S8. Day-folded average locomotor activity profiles of**** Clk ****polyQ alleles at 18 °C and 24 °C.**** Each panel shows the mean daily activity profile for flies carrying one of six Clk polyQ alleles (Q21, Q25, Q29, Q30, Q31, Q33) recorded under a 12:12 light-dark cycle at 24 °C (red) and 18 °C (blue). Separate cohorts of flies were assayed at each temperature. Activity (beam crossings per 30-minute bin) was averaged across recording days within each fly and then across flies within each allele and temperature; shaded envelopes show the standard error of the mean across flies. The x-axis is Zeitgeber time (ZT), with ZT0 corresponding to lights-on and the grey region marking the dark phase (ZT12 to ZT24). Profiles were folded over days 1 to 4 at 24 °C and days 1 to 5 at 18 °C; the number of flies contributing to each profile is given in the panel headers
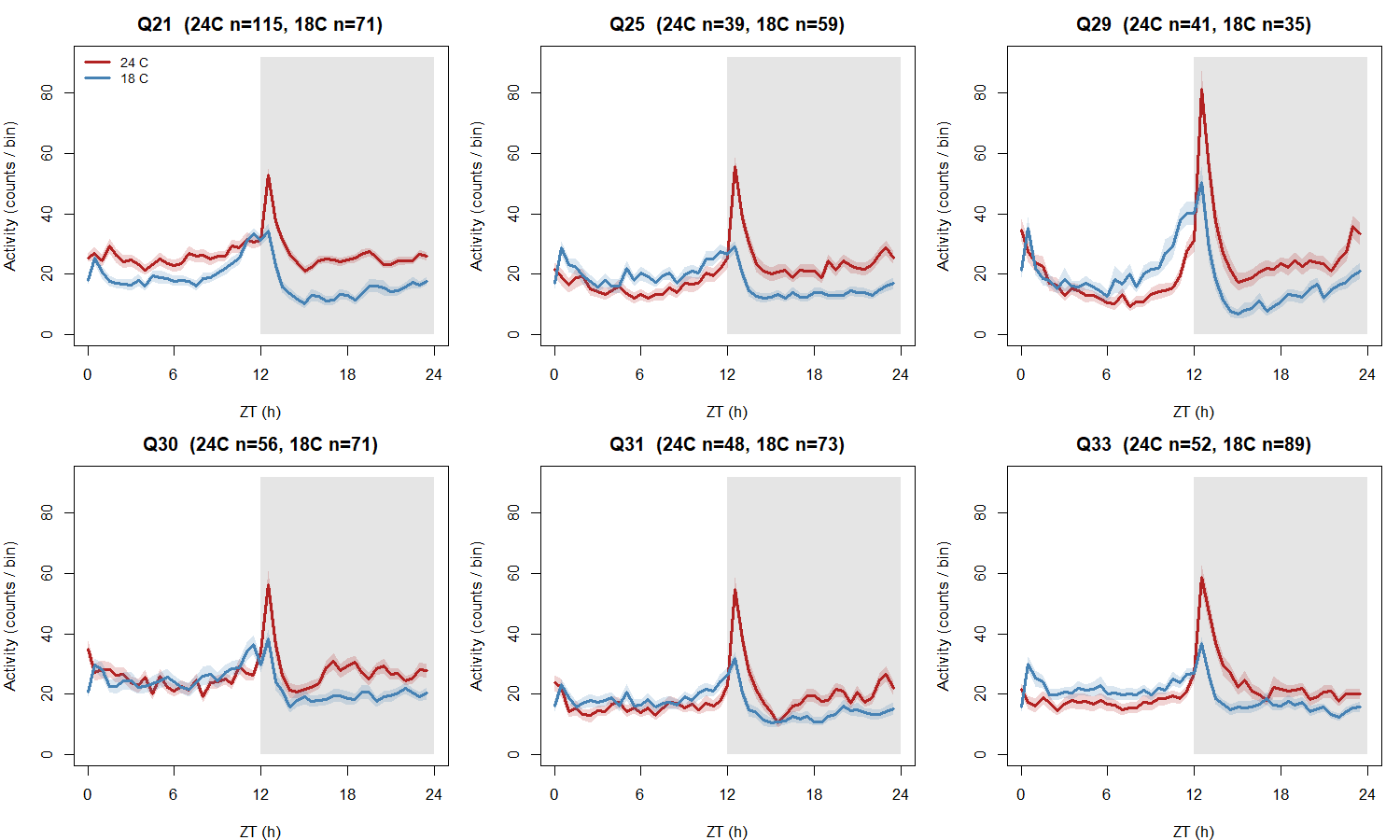
.
